# CFDP1 is required for histone variant H2A.Z deposition by the human SRCAP chromatin remodeling complex

**DOI:** 10.1101/2025.10.30.685587

**Authors:** Naoe Moro, Dandan Yang, Vincent L. Butty, Stuart S. Levine, Craig L. Peterson, Laurie A. Boyer, Shinya Watanabe

## Abstract

Craniofacial Developmental Protein 1 (CFDP1) is a member of the evolutionarily conserved family of Bucentaur (BCNT) proteins and was originally classified as a protein required for cell survival and differentiation during tooth development. Yeast Swc5, a BCNT family member, is an essential subunit of the yeast SWR1C chromatin remodeling complex that catalyzes the deposition of histone variant H2A.Z. Direct connections between CFDP1, H2A.Z deposition, and the mammalian SWR1 homolog, Snf2-Related CREBBP Activator Protein (SRCAP), have not been identified. Here, we perform detailed biochemical reconstitution and characterization of the human SRCAP complex (SRCAP-C). We find that CFDP1 weakly interacts with SRCAP-C in a salt concentration-dependent manner. SRCAP-C purified under a high-salt condition does not co-purify with CFDP1 and is inactive in H2A.Z dimer exchange reaction, but the addition of exogeneous CFDP1 restores the H2A.Z deposition activity of SRCAP-C, demonstrating that CFDP1 is required for H2A.Z dimer exchange by SRCAP-C. We show that CFDP1 stimulates the basal ATPase activity of reconstituted SRCAP-C, suggesting a requirement for CFDP1 in regulating intrinsic catalytic ATPase activity. Consistent with this idea, CFDP1 deficiency in human induced pluripotent stem cells (hiPSCs) leads to a genome-wide reduction of H2A.Z, H3K27me3, and H3K4me3 deposition, accompanied by the upregulation of developmental genes normally marked by these modifications. Taken together, our results provide mechanistic insights into how CFDP1 regulates histone variant H2A.Z deposition by SRCAP-C. Given mutations in the SRCAP gene cause Floating-Harbor syndrome (FHS), a rare, dominant developmental disorder, our study provides an additional link between craniofacial defects and SRCAP-mediated H2A.Z deposition.

## INTRODUCTION

Craniofacial Developmental Protein 1 (CFDP1) is a member of the evolutionarily conserved family of Bucentaur (BCNT) proteins. Mammalian CFDP1 was originally classified as a protein required for cell survival and differentiation during tooth development (*1, 2*). CFDP1 is also involved in craniofacial development in zebrafish (*3*). Yeast Swc5, a BCNT family member, is an essential subunit of the yeast SWR1C chromatin remodeling complex that catalyzes the deposition of histone variant H2A.Z (*4*). However, direct connections between CFDP1, H2A.Z deposition, and the mammalian SWR1 homolog, Snf2-Related CREBBP Activator Protein (SRCAP), have not been identified.

Mutations in the SRCAP gene cause Floating-Harbor syndrome (FHS) (*5–9*). FHS is a rare, dominant developmental disorder characterized by short stature, delayed osseous maturation, delayed speech development, and a dysmorphic facial appearance (*10–12*). All SRCAP mutations in FHS patients are heterozygous truncating alleles, tightly clustered within the final 33^rd^ and 34^th^ exons. Importantly, it was reported that two individuals carrying a 208 kb deletion of a chromosomal region containing the SRCAP gene had no reported phenotype, indicating that SRCAP deletion is haplosufficient and the truncated SRCAP proteins in FHS patients produce a dominant negative effect (*5*).

SRCAP is the catalytic ATPase subunit of the large, multi-subunit SRCAP-C chromatin remodeling complex. SRCAP-C is highly related to the yeast SWR1C remodeler, both structurally and functionally (*13, 14*). SWR1C is a member of the INO80 remodeler subfamily, and it uses the energy of ATP hydrolysis for a unique dimer exchange activity that deposits the histone variant H2A.Z into nucleosomes (*13*). SRCAP is required for H2A.Z deposition in mouse embryonic stem cells (ESCs) and loss of SRCAP leads to upregulation of H3K4me3 and H3K27me3 marked bivalent genes, where H2A.Z is enriched at promoters (*15*). One characteristic feature of INO80 subfamily members is the presence of a large insertion domain between the ATPase lobes, creating a split ATPase domain. This insertion domain is relatively small for the yeast members (247–282 amino acids) but more than 1000 amino acids in length for SRCAP (**Fig. 1A**) (*13*). This insertion domain serves as a docking site for several key subunits of SWR1C-like complexes, including the Rvb1/Rvb2 heterohexameric ring (RUVBL1/RUVBL2 in humans) which acts as a further scaffold for organizing additional subunits. In addition to Rvb1/Rvb2, studies have demonstrated H2A.Z deposition by SWR1C requires several key subunits including Swc2 (mammalian YL1), Swc4 (mammalian DMAP1), Yaf9 (mammalian GAS41), and Swc5 (mammalian CFDP1) (*4, 16*); however, the molecular mechanism by which SRCAP-C catalyzes H2A.Z deposition is largely unknown, mainly due to limited reconstitution experiments as SRCAP-C forms a mega-Dalton multi-protein complex. The contribution of each subunit to the H2A.Z deposition activity of SRCAP-C is also unknown.

**Figure 1.**
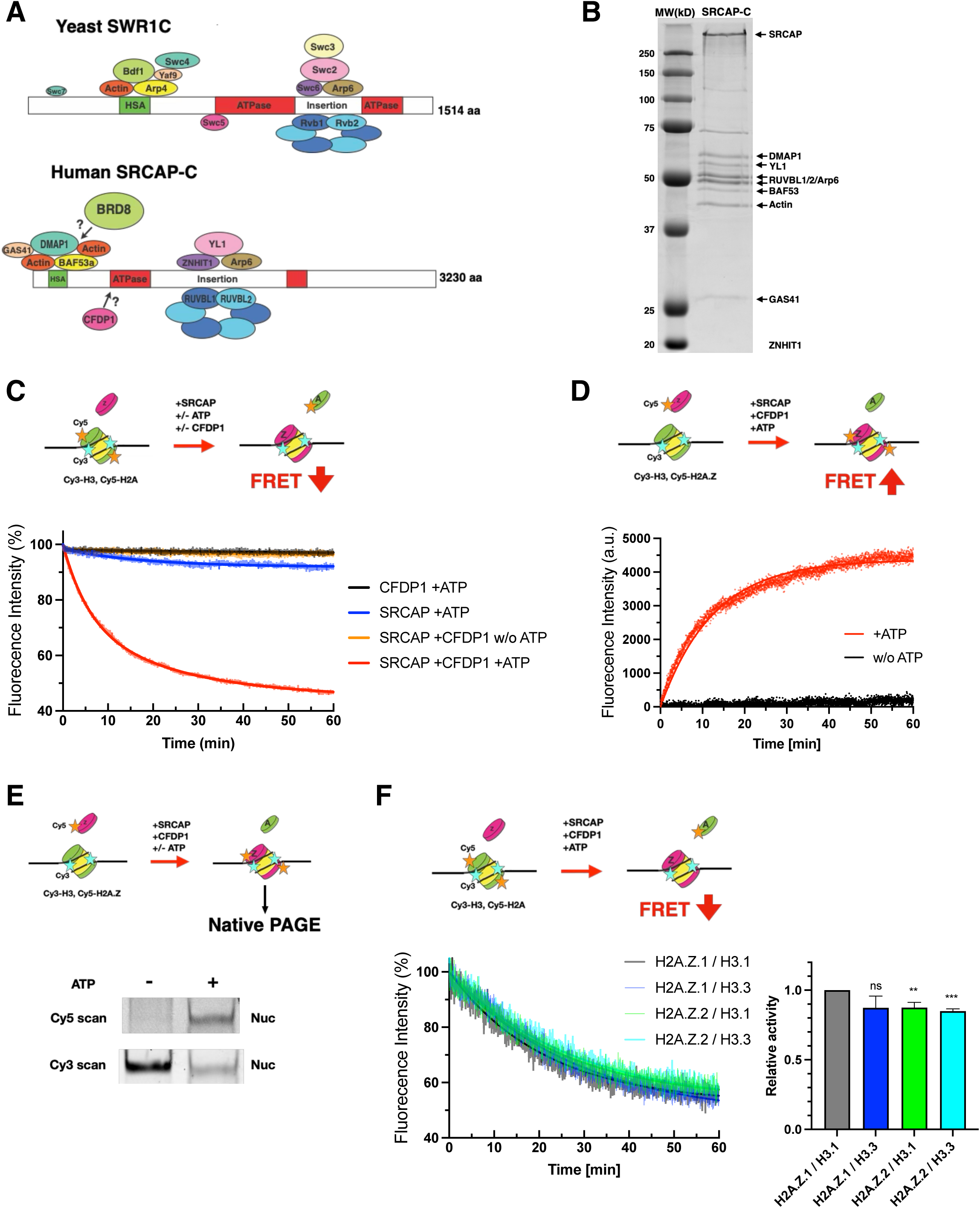
CFDP1 is required for H2A.Z dimer exchange activity by SRCAP-C. (A) Subunit interactions of yeast SWR1C and human SRCAP-C. Subunit interactions of SRCAP-C are predicted based on yeast SWR1C. The homologs are presented in the same colors. (B) Coomassie-staining SDS-PAGE of the purified SRCAP-C. (C) Dimer exchange activity (dimer eviction) of SRCAP-C purified under the high-salt condition. Scheme of FRET-based dimer eviction assay (Top panel). Representative changes in FRET fluorescence intensity over time using a PC1 fluorescent spectrometer with 530 nm excitation and 670 nm emission. (70 µL reaction volume) (Bottom panel). 15 nM SRCAP-C was incubated with 20 nM H3.1-containing mononucleosomes and 150 nM H2A.Z.1/H2B dimers in the presence or absence of 0.5 µM CFDP1 and 2 mM ATP. (D) Dimer exchange activity (dimer deposition) of SRCAP-C. Scheme of FRET-based dimer deposition assay (Top panel). Representative changes in FRET fluorescence intensity over time using a PC1 fluorescent spectrometer (70 µL reaction volume) (Bottom panel). The assays were performed in the same condition as described in Fig. 1C. (E) Gel-based assay for dimer exchange activity (dimer deposition) of SRCAP-C. Scheme of gel-based dimer deposition assay (Top panel). Representative Native-PAGE image of the reactions with Cy3 and Cy5 scans (Bottom panel). The assays were performed in the same condition as described in Fig. 1C. (F) Substrate specificity for histone H2A.Z and H3 variants in SRCAP-C dimer exchange. Scheme of FRET-based dimer eviction assay (Top panel). Representative changes in FRET fluorescence intensity over time using a Tecan plate reader (70 µL reaction volume) (Bottom left panel). The assays were performed in the same condition as described in Fig. 1C. Quantification of dimer exchange activity (Bottom right panel). Each activity was normalized to the activity with H3.1-containing nucleosomes and free H2A.Z.1/H2B dimers. Each error bar represents the standard deviation from four independent experiments using two independent nucleosome/dimer preparations. ***P<0.001, **P<0.01; ns, not significant.

In addition to canonical histones, mammals have a variety of histone variants, including H2A and H3 variants. The variant or combination of variants determines nucleosome stability and function, and variants often show discreet distribution patterns across the genome (*17*). Among them, the histone variant H2A.Z is highly conserved from yeast to human, and it is involved in a wide variety of DNA metabolic processes including gene transcription, DNA repair, and DNA replication (*17*). Two distinct H2A.Z isoforms, H2A.Z.1 and H2A.Z.2 have been identified in the vertebrate genome as products of two non-allelic genes that differ by only three amino acids. H2A.Z.1 knockout studies in mice suggest that the two H2A.Z genes are non-redundant (*18*), and their isoform-specific distributions and functions have been clarified (*17, 19*). The distinct functions of H2A.Z.1 and H2A.Z.2 suggest that each isoform may be deposited through different mechanisms, which may lead to distinct biological consequences. For example, substantial depletion of H2A.Z.1 using a degron tag in mouse ESCs does not affect H2A.Z.2 enrichment at promoters; however, these cells exhibit upregulation of developmental genes and fail to differentiate properly (*19*).

A recent cryo-EM analysis of SRCAP-C provided mechanistic insights into how SRCAP-C binds to a nucleosome, but this reconstituted SRCAP-C with 10 subunits was inactive for H2A.Z dimer exchange activity (*20*), suggesting that a key subunit(s) is missing, precluding studies of the contribution of individual components to its activity. Thus, we reconstituted human SRCAP-C with 12 subunits using a MultiBac baculovirus expression system (*21*). We identify CFDP1 as a previously unidentified SRCAP-C subunit, essential for its H2A.Z dimer exchange activity. Previous studies showed that CFDP1 co-immunoprecipitated with ARP6 and ZNHIT1 subunits of SRCAP-C, but not with SRCAP (*22*), and it was not detected in the highly purified native enzyme complex (*23*). We find that the CFDP1 interaction with SRCAP-C is salt-concentration dependent, and SRCAP-C purified under a high-salt buffer condition does not co-purify with CFDP1 and lacks the H2A.Z deposition activity. We also show that the BCNT domain of CFDP1 is essential for the dimer exchange activity. Our findings also elucidate the key subunits that regulate the ability of SRCAP-C to deposit the histone variant H2A.Z into chromatin. Consistent with the *in vitro* data, CFDP1 is necessary for H2A.Z chromatin deposition and regulation of developmental gene expression in human induced pluripotent stem cells (hiPSCs), similar to prior H2A.Z loss of function studies. Taken together, our results provide mechanistic insights into how CFDP1 regulates histone variant H2A.Z deposition by SRCAP-C and an additional link between craniofacial defects and SRCAP-mediated H2A.Z deposition.

## RESULTS

### CFDP1 is essential for H2A.Z dimer exchange activity by SRCAP-C

To investigate the biochemical properties of SRCAP-C, we reconstituted human SRCAP-C using the MultiBac baculovirus expression system. Based on the similarity of human SRCAP-C and yeast SWR1C **(Fig. 1A)**, we co-expressed Strep-tagged human SRCAP with 11 conserved human proteins (RUVBL1, RUVBL2, YL1, ARP6, ZNHIT1, BRD8, BAF53a, ß-actin, DMAP1, GAS41, and CFDP1). Human SRCAP-C was purified to near homogeneity by Strep purification and glycerol gradient centrifugation. Sodium dodecyl sulfate (SDS)-polyacrylamide gel electrophoresis (PAGE) and mass spectrometry analyses were used to verify subunit identities **(Fig. 1B)**. We conducted quantitative mass spectrometry to measure subunit stoichiometries in SRCAP-C. This analysis confirmed six copies of RUVBL1/RUVBL2 per SRCAP **(Supplementary Table 1)**. These data are consistent with yeast SWR1C that harbors a single heterohexameric ring of Rvb1/Rvb2 (yeast homologs of RUVBL1/RUVBL2) (*24, 25*). When SRCAP-C was purified under high-salt buffer conditions (350 mM NaCl), mass spectrometry detected only a few peptides of CFDP1 and BRD8. Under low-salt buffer conditions (150 mM NaCl), both proteins co-purified with SRCAP-C but at substoichiometric levels **(Supplementary Table 1),** suggesting that CFDP1 and BRD8 interact only weakly with the complex in salt concentration-dependent manner.

To monitor SRCAP-C dimer exchange activity, we performed a FRET-based assay that enables real-time quantitative measurement of the exchange reaction. To evaluate dimer eviction, we labelled histone H2A with Cy5 and histone H3.1 with Cy3 **(Fig. 1C**, **Top)**. Nucleosome assembly brings these fluorophores into close proximity allowing FRET while eviction of H2A during the exchange reaction reduces FRET efficiency. We found that SRCAP-C purified under the high-salt condition showed no detectable dimer exchange activity, whereas SRCAP-C purified under the low-salt condition exhibited mild activity **(Fig. 1C**, **Bottom and Supplementary Fig. 1A)**. Since CFDP1did not co-purify with SRCAP-C in the high-salt condition, we purified recombinant CFDP1 **(Supplementary Fig. 1B)** for subsequent reconstitution experiments. Upon addition of recombinant CFDP1, both the high-salt and low-salt SRCAP-C preparations exhibited robust ATP-dependent dimer exchange activity, whereas CFDP1 alone (as a control) showed no detectable dimer eviction activity **(Fig. 1C**, **Bottom and Supplementary Fig. 1A)**. This dimer exchange activity was also dependent on SRCAP concentration **(Supplementary Fig. 1C)**. Together, these results demonstrate that CFDP1 is an essential subunit for SRCAP-C H2A.Z dimer exchange activity. Thus, all subsequent biochemical experiments were performed using SRCAP-C purified under the high-salt condition in the presence of recombinant CFDP1 to ensure full SRCAP-C activity.

To confirm that the observed reaction by SRCAP-C was due to dimer exchange, we used a FRET-based assay to monitor dimer deposition by SRCAP-C using free, Cy5-labelled H2A.Z.1/H2B dimers and nucleosomes containing Cy3-labelled canonical H3.1 **(Fig. 1D, Top)**. Consequent deposition of H2A.Z.1/H2B dimers into nucleosomes increased FRET efficiency. In the presence of SRCAP-C together with recombinant CFDP1, we observed an ATP-dependent increase in FRET signal indicating that SRCAP-C mediates H2A.Z.1/H2B deposition into nucleosomes **(Fig. 1D, Bottom)**. Together with the dimer eviction assay, these results demonstrate that SRCAP-C catalyzes the H2A.Z dimer exchange reaction, evicting H2A/H2B dimers and incorporating H2A.Z/H2B dimers into the nucleosome.

In addition to the FRET-based assays, we confirmed H2A.Z deposition by SRCAP-C using a gel-based assay, in which free Cy5-labelled H2A.Z.1/H2B dimers were incubated with nucleosomes containing Cy3-labelled H3.1 **(Fig. 1E, Top)**. After incubation with SRCAP-C and recombinant CFDP1 in the presence or absence of ATP, reactions were resolved by native-PAGE and scanned for Cy5 and Cy3 signals. Incorporation of Cy5-labelled H2A.Z.1 into nucleosomes was detected specifically in the presence of ATP **(Fig. 1E)**. Notably, the Cy3 signal from H3.1 decreased in the presence of ATP, consistent with FRET from Cy3 on H3.1 to Cy5 on the incorporated H2A.Z. Taken together, both FRET-based and gel-based assays confirmed the H2A.Z dimer exchange activity of SRCAP-C indicating that CFDP1 (Swc5 in yeast) is indeed necessary for both human and yeast complexes.

### SRCAP-C has no substrate preference for several histone variants

Histone H3 variant H3.3 is deposited in transcriptionally active chromatin and is mostly co-localized with H2A.Z in active gene promoters, enhancers, and insulator regions (*17*). Nucleosomes harboring both H2A.Z and H3.3 are unstable and easily disassembled suggesting that they facilitate access to these sites for transcription factors (*26*). Although the deposition mechanism of co-localized H2A.Z and H3.3 is unclear, one possible explanation is that SRCAP-C preferentially deposits H2A.Z into H3.3-containing nucleosomes over H3.1-containing nucleosomes. To test this idea, we compared histone H3.1-containing and H3.3-containing nucleosomes in the dimer exchange reaction by SRCAP-C. The FRET-based dimer exchange assay showed no significant difference in the H2A.Z exchange rate between histone H3.1 and H3.3 **(Fig. 1F)**, suggesting that SRCAP-C has no substrate preference between these histone H3 variants. We also compared free H2A.Z.1/H2B dimers and H2A.Z.2/H2B dimers in the dimer exchange reaction by SRCAP-C. The FRET-based assay showed H2A.Z.1 was significant, but only modest preference over H2A.Z.2 on both H3.1-containing and H3.3-containing nucleosomes **(Fig. 1F)**. Similar results were obtained for the two independent preparations of either histone H3 or H2A.Z histone variants. These results suggest that SRCAP-C has no substrate preference among these H2A.Z and H3 variants in the dimer exchange reaction.

### CFDP1 stimulates basal ATPase activity of SRCAP-C

To understand the mechanism by which CFDP1 mediates H2A.Z deposition by SRCAP-C, we investigated whether CFDP1 is necessary for the ATPase activity of the complex. ATPase activity was monitored by fluorescent-labelled phosphate binding protein (MDCC-PBP), which binds to phosphate released from ATP hydrolysis leading to an increase in fluorescence (*27*). First, we measured the basal ATPase activity of SRCAP-C in the absence of DNA or a nucleosomal substrate and in the presence of varying concentrations of CFDP1. The ATPase activity of SRCAP-C was stimulated by CFDP1 in a concentration-dependent manner (**Fig. 2A**). Note that no fluorescence change was observed in the presence of CFDP1 alone (**Fig. 2A, Right**). These results suggest that CFDP1 activates the intrinsic catalytic ATPase activity of SRCAP-C.

**Figure 2.**
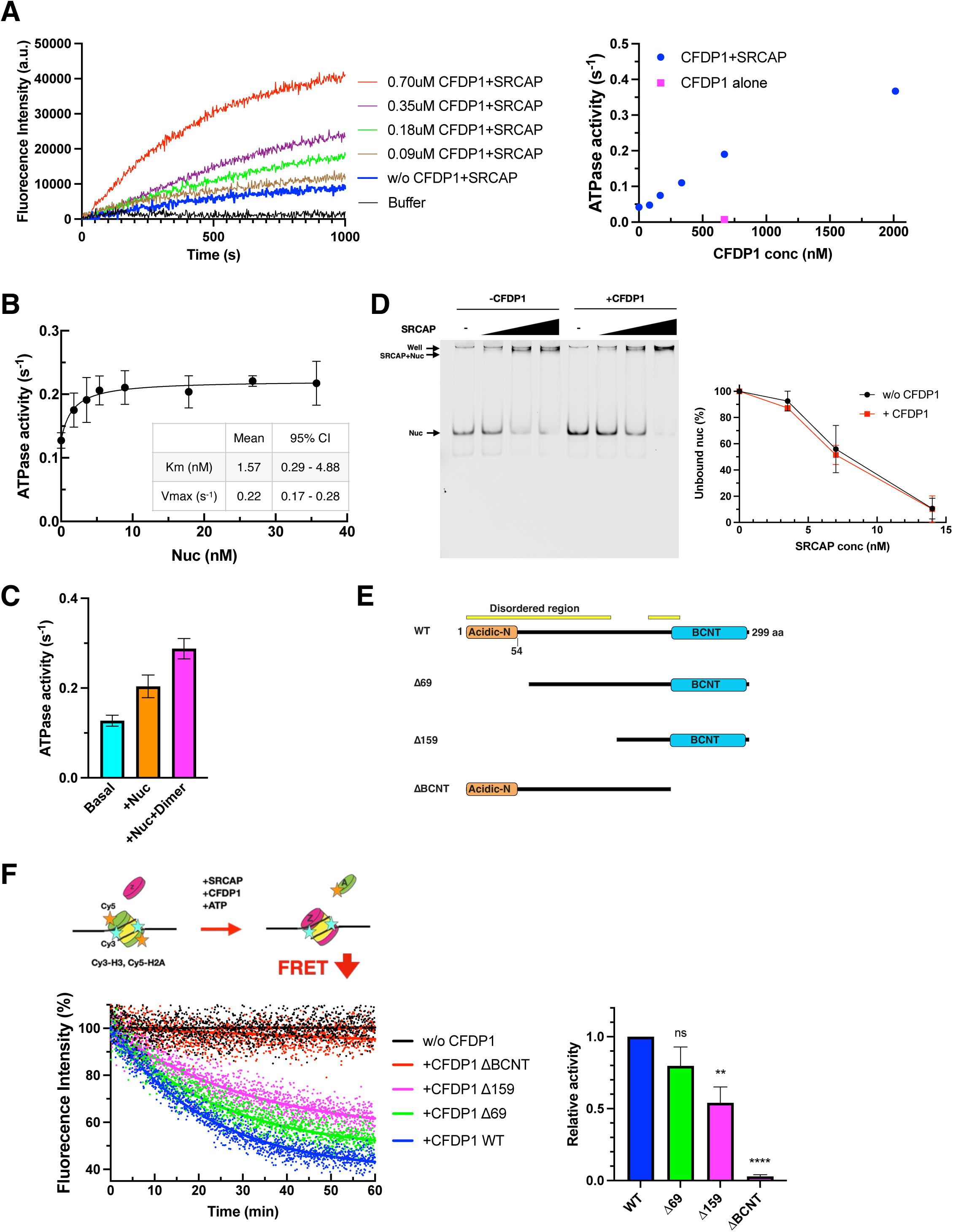
CFPD1 activates the ATPase activity of SRCAP-C. (A) CFDP1-concentration dependence of the basal ATPase activity of SRCAP-C. Representative fluorescence changes of MDCC-PBP over time upon ATP addition in various CFDP1 concentrations (Left panel). The ATPase activity of SRCAP-C at each CFDP1 concentration (Right panel). Note that CFDP1 alone showed no fluorescence change. Data are presented as means from two independent experiments using two independent CFDP1 preparations. (B) Nucleosome-concentration dependence of the ATPase activity of SRCAP-C in the presence of 0.35 µM CFDP1. The ATPase activity was measured using MDCC-PBP. Each error bar represents the standard deviation from three independent experiments. Km and Vmax were calculated from the Michaelis-Menten kinetics equation along with 95% confidence intervals (CI). (C) The ATPase activity of SRCAP-C in the presence of 20 nM nucleosomes, 150 nM free H2A.Z.1/H2B dimers, and 0.35 µM CFDP1. Each error bar represents the standard deviation from three independent experiments. (D) Representative native gel of EMSA for the nucleosome binding of SRCAP-C in the presence or absence of CFDP1 (Left panel). 7 nM mononucleosomes were incubated with SRCAP-C (3.5, 7, and 14 nM) in the presence or absence of 1 µM CFDP1 and subjected to 4% native-PAGE. Quantification of unbound nucleosome fractions (Right panel). Each error bar represents the standard deviation from three independent experiments. (E) Domain structure of CFDP1 derivatives. (F) Dimer exchange activity of SRCAP-C with CFDP1 derivatives. Scheme of FRET-based dimer eviction assay (Top panel). Representative changes in FRET fluorescence intensity over time using a Tecan plate reader (20 µL reaction volume) (Bottom left panel). The assays were performed as described in Fig. 1C. Quantification of dimer exchange activity (Bottom right panel). Each activity was normalized to the wild-type CFDP1 activity. Each error bar represents the standard deviation from three independent experiments. \*\*\*\**P*<0.0001, **P<0.01; ns, not significant.

The ATPase activity of yeast SWR1C is stimulated by H2A-containing nucleosomes and further stimulated by free H2A.Z/H2B dimers (*28*). Similarly, in the presence of additional CFDP1, the ATPase activity of SRCAP-C was stimulated by H2A-containing nucleosomes in a concentration-dependent manner **(Fig. 2B)** and further stimulated by free H2A.Z/H2B dimers **(Fig. 2C).**

Next, we investigated the effect of CFDP1 on nucleosome binding of SRCAP-C. Electrophoretic mobility shift assay (EMSA) indicated that CFDP1 neither binds to nucleosomes (**Supplementary Fig. 2**) nor affects the binding of SRCAP-C to nucleosomes (**Fig. 2D**), suggesting that CFDP1 interaction with SRCAP-C influences its catalytic activity.

### The BCNT domain of CFDP1 is required for SRCAP-C-mediated dimer exchange

In yeast, the conserved BCNT domain of Swc5, the CFDP1 homolog, is required for SWR1C mediated H2A.Z exchange, whereas deletion of its conserved acidic N-terminal domain leads to a partial loss of exchange activity (*29*). To test if these domain functions are conserved in human CFDP1, we expressed and purified truncation derivatives of CFDP1 from insect cells **(Fig. 2E**) and assessed their activity in dimer exchange reactions. Loss of the acidic N-terminal domain (Δ69) had a mild effect on the dimer exchange activity of SRCAP-C, but deletion of the BCNT domain (ΔBCNT) resulted in a complete loss of the dimer exchange activity **(Fig. 2F**), consistent with yeast Swc5. The requirement for the BCNT domain in dimer exchange activity is consistent with its role as a chromatin interaction and anchoring module in yeast (*30*). While previous studies demonstrated that the disordered N-terminal region of Swc5 (1-147aa) was essential for dimer exchange activity (*29*), deletion of the same disordered region of CFDP1 (Δ159) only led to a partial loss of the dimer exchange activity **(Fig. 2F**). These data highlight the essential role for the BCNT domain in the mammalian complex.

### YL1 and DMAP1 subunits are essential for dimer exchange by SRCAP-C

To further understand the mechanism of SRCAP-C-mediated dimer exchange, we assessed the requirement of each subunit for dimer exchange activity by eliminating each subunit from our reconstitution system (**Fig. 3A**). Removal of either YL1 or DMAP1 resulted in a complete loss of dimer exchange activity, consistent with previous analyses of the yeast homologs Swc2 and Swc4, respectively (*4, 16*). Although the deletion of YL1 did not impact the integrity of the complex, the removal of DMAP1 resulted in loss of BAF53a and GAS41 from the complex. Likewise, loss of GAS41, which contains a YEATS domain that recognizes a variety of histone modifications, did not disrupt the integrity of the complex but resulting in a partial loss of the dimer exchange activity. These results are consistent with studies of the yeast homolog, Yaf9 (*4, 16*). By contrast, removal of either BAF53a or BRD8 did not affect either the integrity of SRCAP-C or the dimer exchange activity. Similarly, the Brd1 subunit (yeast BRD8 homolog) of SWR1C is not required for H2A.Z deposition, though the yeast homolog of BAF53a, Arp4, is essential for SWR1C activity. However, loss of Arp4 leads to partial loss of additional SWR1C subunits, confounding these functional studies (*16*). Together, these data demonstrate that SRCAP-C and SWR1C rely on a similar set of subunits for optimal H2A.Z deposition activity.

**Figure 3.**
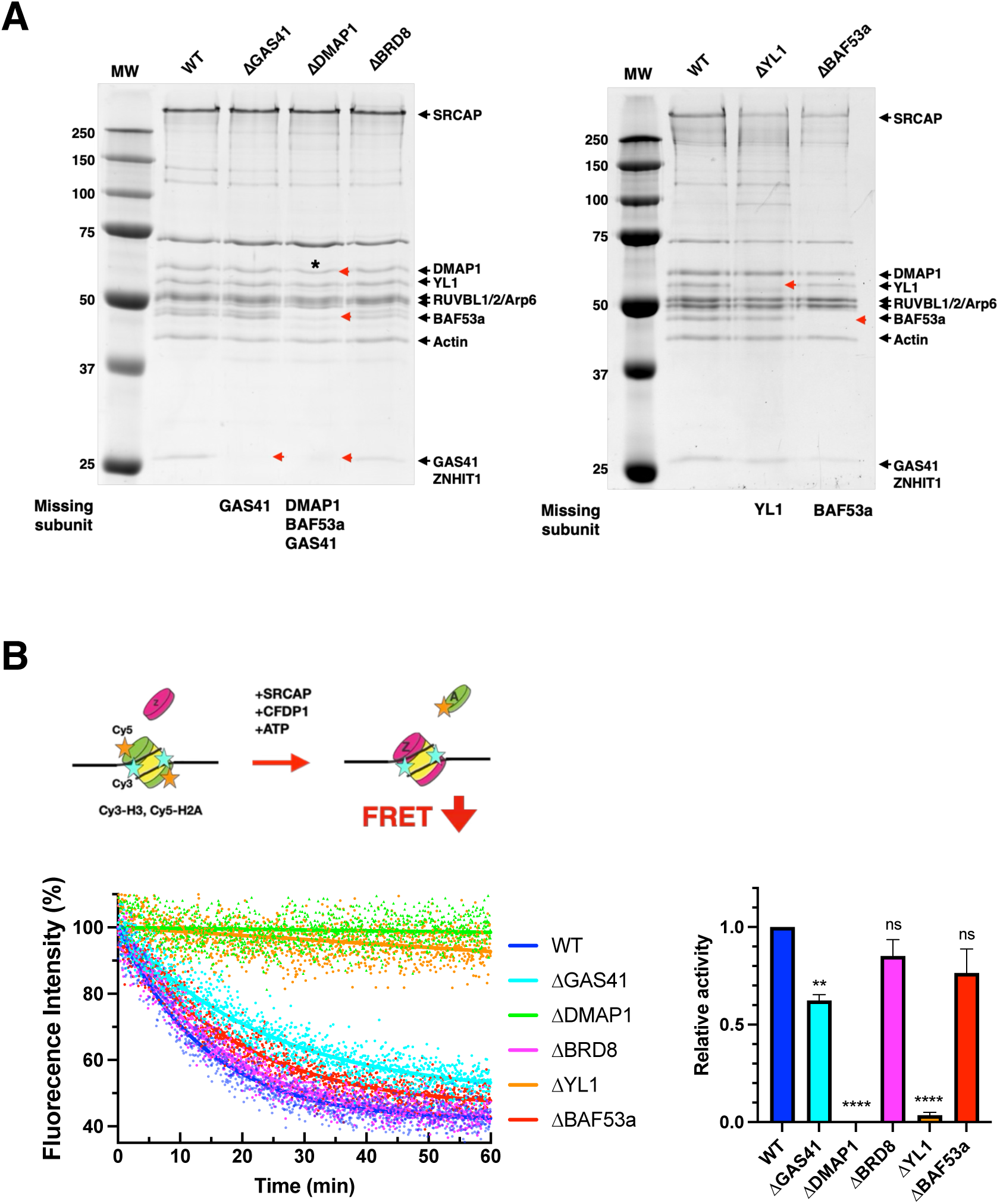
YL1 and DMAP1 subunits are essential for SRCAP dimer exchange. (A) Coomassie-staining SDS-PAGEs of SRCAP-C lacking an indicated subunit. Red arrows, missing subunits. Asterisk, contaminant proteins comigrated with DMAP1. (B) Dimer exchange activity of SRCAP-C lacking an indicated subunit. Scheme of FRET-based dimer eviction assay (Top panel). Representative changes in FRET fluorescence intensity over time using a Tecan plate reader (20 µL reaction volume) (Bottom left panel). The assays were performed as described in Fig. 1C. Quantification of dimer exchange activity (Bottom right panel). Each activity was normalized to the wild-type SRCAP-C activity. Each error bar represents the standard deviation from three independent experiments. \*\*\*\**P*<0.0001, **P<0.01; ns, not significant.

### CFDP1 is necessary for proper H2A.Z deposition in human iPSCs

SRCAP-C has been proposed as the key chromatin remodeler responsible for H2A.Z incorporation, and proper H2A.Z deposition is critical for proper induction of developmental genes (*31, 32*). This balance is essential for both self-renewal and the differentiation potential of stem cells. Given our finding that CFDP1 is required for the H2A.Z dimer exchange activity of SRCAP-C *in vitro*, we next examined the role of CFDP1 in H2A.Z deposition in human iPSCs. To assess global H2A.Z chromatin incorporation, we performed siRNA-mediated knockdown of CFDP1 or SRCAP and observed a robust reduction of CFDP1 and SRCAP at both the mRNA and protein levels (**Supplementary Fig. 3A-D**). Depletion of either CFDP1 (si*CFDP1*) or SRCAP (si*SRCAP*) significantly reduced H2A.Z levels in chromatin fractions extracted with both low-salt (300 mM) and high-salt (500 mM) buffers (**Fig. 4A-B**) compared with control (siScramble). Notably, total cellular levels of H2A.Z remained unchanged across all conditions (**Supplementary Fig. 3E**), indicating that CFDP1 is required for chromatin incorporation of H2A.Z, rather than regulating its overall expression.

**Figure 4.**
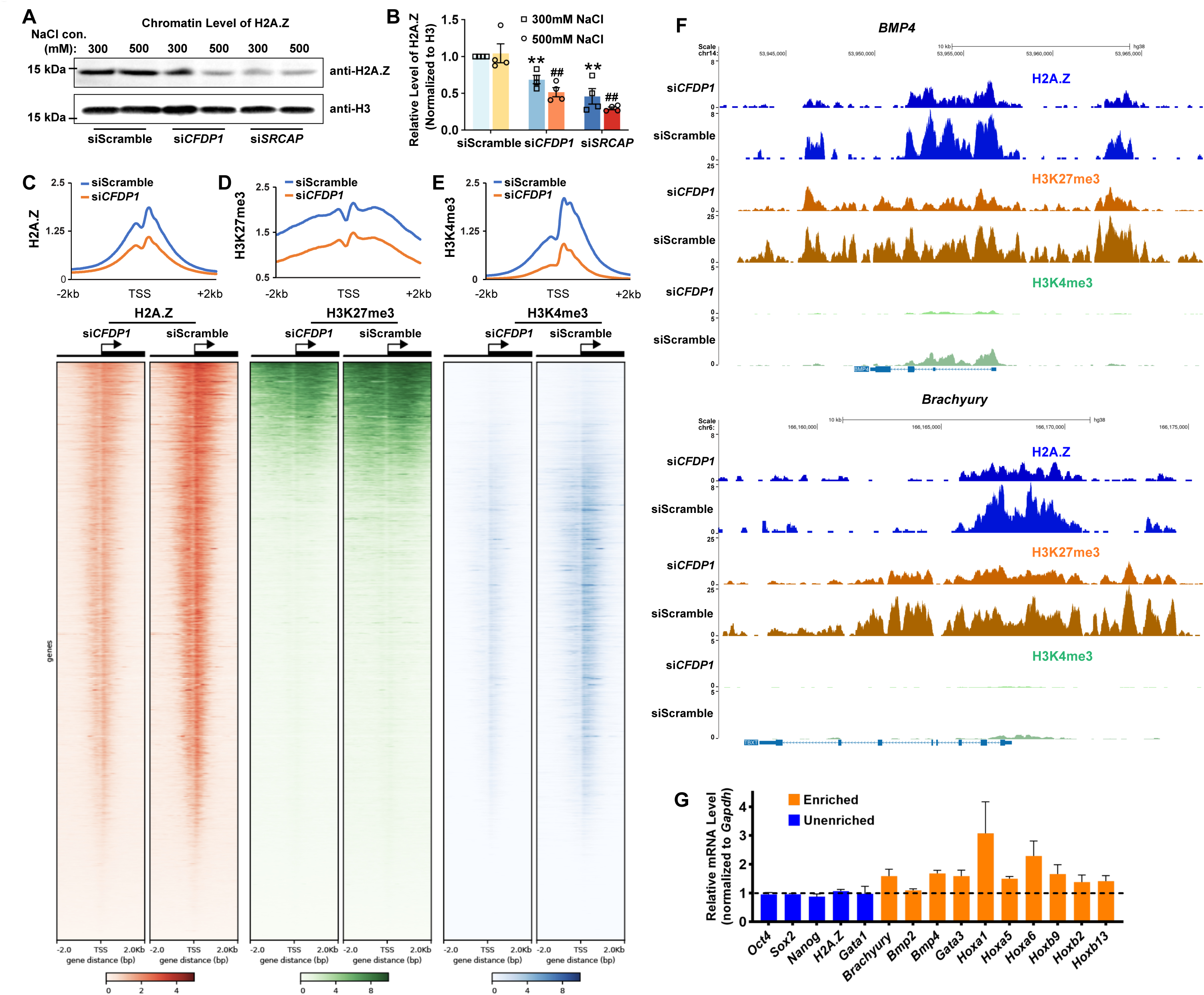
CFDP1 deficiency disrupts H2A.Z deposition into chromatin in the human iPSCs. (A-B) Representative western blot image (A) and quantification of chromatin-associated H2A.Z levels (B) show that siRNA-mediated knockdown of *CFDP1* (si*CFDP1*) or *SRCAP* (si*SRCAP*) reduces H2A.Z incorporation into chromatin. Histones were extracted using the indicated salt concentrations and analyzed by SDS-PAGE. Histone H3 was used as a loading control. *N*=4; error bars represent 2 standard deviations from the mean. \*\**P* < 0.01 versus scramble siRNA (siScramble) with 300 mM NaCl; #*P* < 0.05, ##*P* < 0.01 versus siScramble with 500 mM NaCl. (C-E) Average enrichment profiles and corresponding density maps of H2A.Z (C), H3K27me3 (D) and H3K4me3 (E) within ±2 kb of the transcription start sites (TSSs) reveal a global reduction of these marks in si*CFDP1* cells compared to controls. In the density maps, genes are ranked from highest to lowest based on signal intensity of H2A.Z, with enrichment plotted across a ±2 kb window relative to the TSS. (F) Genome browser tracks of CUT&RUN reads showing the distribution of H2A.Z and H3K27me3 across the *BMP4* locus, a representative developmental gene marked by both modifications. (G) qRT-PCR analysis indicates that H2A.Z target genes (orange) are de-repressed in si*CFDP1* hiPSCs. Data were normalized to *GAPDH* and are presented relative to expression levels in siScramble cells. *N*=4; error bars represent 2 standard deviations from the mean.

In pluripotent stem cells, H2A.Z is broadly deposited near transcription start sites (TSSs), where it regulates the transcriptional activity of developmental gene expression programs (*19, 33*). This regulation occurs in coordination with other chromatin modifications such as H3K4me3 and H3K27me3 and is essential for proper differentiation (*33, 34*). To investigate the impact of CFDP1 on these chromatin features, we performed CUT&RUN to profile the genome-wide distribution of H2A.Z, H3K4me3, and H3K27me3 in human iPSCs following CFDP1 knockdown. As shown in the average density profiles (**Fig. 4C** and **Supplementary Fig. 4A**, **Top**), the characteristic bimodal enrichment of H2A.Z around TSSs was markedly diminished in si*CFDP1* cells compared to the control. This decrease extended across genes with both low and high levels of H2A.Z enrichment (**Fig. 4C** and **Supplementary Fig. 4A**, **Bottom**), indicating a global depletion. Concomitantly, siCFDP1 hiPSCs exhibited a noticeably flatter average profile and significantly reduced overall signal of H3K27me3 and H3K4me3 compared to the control (**Fig. 4D-E** and **Supplementary Fig. 4B-C**). These data are consistent with previous findings that H2A.Z is necessary for maintenance of bivalent chromatin marked by both H3K4me3 and H3K27me3 (*33, 34*). Specifically, we observed a decrease of both modifications at key developmental loci such as *BMP4 and Brachyury* in si*CFDP1* cells compared to controls (**Fig. 4F)**. To test if CFDP1 depletion influences gene expression, we performed qRT-PCR analysis of genes known to be co-enriched for H2A.Z and bivalent chromatin marks. Notably, these genes showed a consistent trend toward de-repression in CFDP1-deficient cells, while genes lacking these chromatin marks exhibited no significant changes (**Fig. 4G)**. Together, our data provide compelling evidence that CFDP1 is essential for SRCAP-mediated H2A.Z deposition and maintaining proper chromatin structure including H3K4me3 and H3K27me3, necessary for proper transcriptional regulation of key developmental genes in human iPSCs.

## DISCUSSION

CFDP1 was originally identified as a protein required for tooth development in mammals. However, a direct connection between CFDP1 and H2A.Z deposition has not been identified. Histone variant H2A.Z is highly conserved from yeast to human and plays crucial roles in transcription, DNA replication, and repair. Furthermore, upregulation of H2A.Z is linked to cancer and cardiac hypertrophy (*35, 36*). Although the mechanism of H2A.Z deposition by the yeast SWR1C remodeler has been well studied, how H2A.Z deposition is regulated in mammalian cells remains to be fully elucidated. Mammalian cells contain two remodeling enzymes related to yeast SWR1C – Tip60/p400 and SRCAP-C. Depletion of the ATPase subunit from either remodeler in mammalian cells leads to depletion of H2A.Z from genomic sites, indicating that they both contribute to H2A.Z deposition *in vivo* (*31, 32, 37, 38*). Here, we reconstituted the human SRCAP-C remodeling enzyme with CFPD1 and other 11 recombinant subunits and demonstrated robust H2A.Z exchange activity in the presence of CFDP1. Analysis of SRCAP-C subcomplexes shows that the YL1, DMAP1, and CFDP1 subunits are essential for ATP-dependent H2A.Z deposition. Notably, CFDP1 had not been previously identified as a critical subunit of SRCAP-C. We find that while the CFDP1 subunit does not appear to impact nucleosome binding, it is required to stimulate the basal ATPase activity of SRCAP-C in the reconstituted complex. Depletion of CFDP1 from human iPSCs leads to a global depletion of H2A.Z, similarly to loss of the SRCAP ATPase (*15*).

A recent study reported the reconstitution of human SRCAP-C with only 10 recombinant subunits and H2A.Z dimer exchange activity was not detected (*20*). Notably, their reconstituted complexes lacked BRD8 and CFDP1, orthologs of the yeast Brd1 and Swc5 subunits of SWR1C, respectively. Previous mass spectrometry analysis of native SRCAP-C purified from HeLa cells did not reveal peptides for CFDP1 (*23*), and immunoprecipitation studies with CFDP1 failed to detect an interaction between CFDP1 and the SRCAP ATPase (*22*). The yeast ortholog of CFDP1, Swc5, is essential for H2A.Z deposition by SWR1C, as it promotes both nucleosome binding and ATPase activity (*29, 39*). Thus, it was surprising that CFDP1 was not associated with SRCAP-C. Our study indicates that CFDP1 does associate with SRCAP-C, but the interaction is salt-sensitive. In our low salt isolation conditions, a substoichiometric level of CFDP1 is associated with SRCAP-C, and this enzyme preparation has measurable H2A.Z deposition activity. In contrast, purification in standard, high salt buffer eliminated both CFDP1 association and H2A.Z deposition activity. These results contrast to the case of yeast Swc5, which is a salt-stable subunit of SWR1C. Our observation of a weak interaction between CFDP1 and SRCAP-C may reflect more dynamic regulation of SRCAP-C activity by CFDP1 in mammalian cells.

The initial cryoEM structural model of a yeast SWR1C-nucleosome complex failed to visualize the key Swc5 subunit, presumably due to flexibility (*25*). Notably, this structural model is highly related to that of SRCAP-C-nucleosome models that lack the CFDP1 ortholog (*20*). A more recent cryoEM structural model of SWR1C bound to a hexasome did visualize the conserved, C-terminal BCNT domain of Swc5 (*30*), which is known to be essential for Swc5 function in vivo and for SWR1C activity *in vitro* (*29*). In this structural model, the BCNT domain stretches out in contact with unwrapped DNA and histones, suggesting that Swc5 stabilizes an intermediate nucleosomal substrate in the dimer exchange reaction (*30*). The cryoEM model also showed that the Swc5 BCNT domain wraps across the ATPase domain of the Swr1 subunit, providing structural support for ATPase stimulation. We found that the BCNT domain of CFDP1 is also essential for the dimer exchange activity of SRCAP-C, and given the conservation of the BCNT domain between yeast and human, it is likely that the BCNT domain of CFDP1 also interacts with the ATPase domain of the SRCAP subunit, activates the ATPase activity, and mediates the dimer exchange reaction by stabilizing an unwrapped intermediate nucleosomal substrate.

Co-localization of H2A.Z and histone H3.3 has been observed in various genome loci such as active promoters, enhancers, and insulator regions (*26*). Although the mechanisms that lead to this colocalization are unclear, we tested the possibility that SRCAP-C might preferentially deposit H2A.Z into H3.3-containing nucleosomes over H3.1-containing nucleosomes. However, our dimer exchange assays indicated that SRCAP-C shows no detectable substrate preference between H3.1 and H3.3, suggesting that the recruitment of SRCAP-C to H3.3-containing nucleosomes may be the primary mechanism that leads to co-localization of H2A.Z and H3.3. Consistent with this model, a recent study showed that the HIRA complex, a histone H3.3 chaperone, interacts with SRCAP-C, supporting a model in which SRCAP-C is recruited to H3.3-containing nucleosomes through the HIRA complex (*40*).

Interestingly, a recent study demonstrated that H2A.Z.2 knockdown and overexpression mimic and rescue the FHS phenotype, respectively (*41*), suggesting a functional connection between H2A.Z.2 and FHS, yet the underlying mechanism is unclear. Since our study showed that SRCAP-C has no significant substrate preference between H2A.Z.1 and H2A.Z.2, it is possible that SRCAP-C FHS mutant may have a different substrate preference or other mechanisms may exist for histone variant specific deposition or removal including coordination with Tip60/p400 and specific histone chaperones. Thus, our biochemical and functional analyses of SRCAP-C provide invaluable insights for understanding the roles of SRCAP and histone variant exchange in development and complex disease.

## METHODS

### Insect cell culture

Sf9 and HighFive cells (Thermo Fisher Scientific, Waltham, MA, USA) were used for baculovirus production and recombinant protein expression, and grown in ESF921 media (Expression Systems, Davis, CA, USA) at 27 °C.

### Expression and purification of human SRCAP complex

Human SRCAP subunit genes were cloned and expressed using the MultiBac baculovirus expression system (Geneva Biotech, Geneva, Switzerland). Genes coding for WT SRCAP (residue 1–3230 with an N-terminal 3× FLAG-tag and a C-terminal twin Strep-tag), ARP6, β-Actin, BAF53a, BRD8, CFDP1, DMAP1, GAS41, RUVBL1, RUVBL2, YL1, and ZNHIT1 were synthesized with codon optimization for insect cells (Genewiz, Cambridge, MA, USA).

HighFive cells were co-infected with two viruses expressing all subunits, and cultured for 72 h at 27 °C. Cells were harvested and washed with phosphate-buffered saline and then lysed by sonication in Lysis buffer (150 mM [low-salt] or 350 mM [high-salt] NaCl, 20 mM HEPES [pH 7.5], 10% glycerol, 0.1% Tween, 1 mM MgCl_2_, 50 µM ZnCl_2_, 1 mM DTT, 1 mM phenylmethylsulphonyl fluoride, 1 mM benzamidine, 2 µg/mL Leupeptin, 2 µg/mL Pepstatin A, 2 µg/mL Chymostatin, and 5 µL benzonase nuclease [Sigma-Aldrich, St. Louis, MO, USA]). After centrifugation at 40,000 *g* and 4 °C for 30 min, the supernatant was loaded onto a StrepTactin HP column (Cytiva, Marlborough, MA, USA). The column was washed with Lysis buffer and buffer A (150 mM NaCl, 20 mM HEPES [pH 7.5], 10% glycerol, 1 mM MgCl_2_, and 1 mM DTT) and eluted with buffer A plus 10 mM desthiobiotin (Sigma-Aldrich). The eluted proteins were subjected to 15–40% glycerol gradient centrifugation. The peak fractions were concentrated in buffer A and flash-frozen in liquid nitrogen. Protein concentration was determined by the densitometry analysis of SDS-PAGE using BSA as a standard. Subunit compositions were confirmed by SDS-PAGE and mass spectrometry. Note that we added benzonase nuclease to the lysis and wash buffers in both salt buffer conditions to avoid protein interactions through contaminated DNA.

To purify recombinant CFDP1 protein, CFDP1 with a C-terminal 3× FLAG-tag was expressed in Sf9 cells. Harvested cells were lysed by sonication in Lysis buffer with 350 mM NaCl. Cleared lysate was incubated with Anti-FLAG M2 affinity gel (Sigma-Aldrich) at 4 °C for 1 hour, washed with Lysis buffer and buffer A, and eluted with buffer A containing 0.1 mg/ml 3× FLAG peptide (GenScript, Piscataway, NJ, USA). The peak fractions were concentrated in buffer A and flash-frozen in liquid nitrogen.

### Nucleosome preparation

Recombinant human histones were expressed in *Escherichia coli* cells and purified as previously described (*42, 43*). In brief, expressed histones were purified as inclusion bodies, solubilized in unfolding buffer (7 M guanidinium hydrochloride, 20 mM Tris-HCl [pH 7.5], and 10 mM DTT), and dialyzed against urea dialysis buffer (7 M urea, 10 mM Tris-HCl [pH 8.0], 0.1 M NaCl, 1 mM EDTA, 0.2 mM phenylmethylsulphonyl fluoride, and 5 mM 2-mercaptoethanol). Samples were injected into tandemly connected Q Sepharose and SP Sepharose columns, and eluted from SP Sepharose by a linear salt gradient. Histone fractions were dialyzed against water with 0.2 mM phenylmethylsulphonyl fluoride and 5 mM 2-mercaptoethanol, and lyophilized. Histones H2A (T120C), H2A.Z.1 (G122C), and H2A.Z.2 (G122C) were labelled with Cy5; histones H3.1 (G33C, C96S, C110A) and H3.3 (G33C, C110A) were labelled with Cy3, as previously described (*44*). The four histones (H2A, H2B, H3, and H4) for octamers and the two histones (H2A and H2B or H2A.Z and H2B) for dimers were mixed in equimolar ratios in unfolding buffer, dialyzed against refolding buffer (2 M NaCl, 10 mM Tris-HCl [pH 7.5], 1 mM EDTA, and 5 mM 2-mercaptoethanol) and purified through a Superdex-200 column. Nucleosomes were reconstituted by mixing octamers with DNA at a 1:1 ratio in HI buffer (2 M NaCl, 10 mM Tris-HCl [pH 7.5], and 5 mM 2-mercaptoethanol), and dialyzed against a linear salt gradient buffer from HI to LO buffer (50 mM NaCl, 10 mM Tris-HCl [pH 7.5], and 5 mM 2-mercaptoethanol) for 20 h.

### FRET-based dimer exchange assays

For dimer eviction assays, mononucleosomes with Cy5-labelled H2A and Cy3-labelled H3 were reconstituted by salt dialysis onto a 250 bp DNA fragment containing the 601 nucleosome-positioning sequence in the center of the DNA fragment. Mononucleosomes (20 nM) were incubated with SRCAP-C, free histone H2A.Z/H2B dimers (150 nM) and 2 mM ATP in buffer B (60 mM NaCl, 25 mM HEPES [pH 7.5], 5 mM MgCl_2_, and 1 mM DTT) at 37 °C for 60 min. The FRET signal was monitored using a PC1 fluorescent spectrometer (ISS, Champaign, IL, USA) or a Tecan Spark microplate reader (Tecan, Switzerland) with excitation at 530 nm and emission at 670 nm. Dimer exchange activity was obtained from an initial linear rate. Note that since the reaction volumes were 70 uL for a PC1, and 70 uL (96 wells) or 20 uL (384 wells) for a Tecan, signal to noise ratios were different among these experiments.

For dimer deposition assays, mononucleosomes containing Cy3-labelled H3 were reconstituted by salt dialysis onto the 250 bp DNA fragment. Mononucleosomes (20 nM) were incubated with SRCAP-C, free Cy5-lebelled histone H2A.Z/H2B dimers (150 nM) and 2 mM ATP in buffer B (60 mM NaCl, 25 mM HEPES [pH 7.5], 5 mM MgCl_2_, and 1 mM DTT) at 37 °C for 60 min. The FRET signal was monitored using a PC1 fluorescent spectrometer with excitation at 530 nm and emission at 670 nm.

For gel-based dimer exchange assays, the assays were performed as described in the FRET-based dimer deposition assays except, after 60 min incubation, the reactions were quenched with 5% glycerol and 0.1 mg/ml salmon sperm DNA, incubated for 5 min at 37 °C, and resolved on 5% Native-PAGE in 0.5 X TBE. Gels were scanned with Cy3 and Cy5 using a Typhoon biomolecular imager (GE Healthcare, Chicago, IL, USA).

### ATPase assays

ATPase activity of SRCAP-C was measured as previously described (*27, 45*) using 7-diethylamino-3-((((2-maleimidyl)-ethyl)amino)carbonyl)coumarin-labeled phosphate-binding protein (MDCC-PBP, Thermo Fisher Scientific). Fluorescent changes in MDCC-PBP upon phosphate binding were monitored using a Tecan Spark microplate reader with excitation at 430 nm and emission at 460 nm (70 uL reaction volume).

All solutions were preincubated with 0.5 µM MDCC-PBP and a “Pi mop” (0.01 units/mL purine nucleoside phosphorylase and 0.2 mM 7-methylguanosine) to remove free inorganic phosphate. The assays were performed at 37 °C with 0.5 nM SRCAP-C and 0.1 mM ATP in buffer B. ATPase activity was obtained from an initial linear rate subtracted from background signal (without SRCAP-C).

### Gel-shift assays for nucleosome binding

Mononucleosomes (7 nM) were incubated with remodeling enzymes in buffer B with 5% glycerol for 15 min at room temperature. The reactions were resolved on 4% Native-PAGE in 0.25× TBE. Gels were scanned using a Typhoon biomolecular imager.

### Cell culture

The human induced pluripotent stem cell (hiPSC) line (Penn0141-37-3) was generously provided by the University of Pennsylvania under a material transfer agreement. hiPSCs were maintained on Matrigel-coated plates (Corning, Cat# 354277) in mTeSR Plus medium (STEMCELL Technologies, Cat# 100-0276), with medium changes every two days. Cells were passaged using ReLeSR (STEMCELL Technologies, Cat# 100-0483) according to the manufacturer’s protocol.

### siRNA-mediated knockdown

For siRNA-mediated gene knockdown, hiPSCs were first dissociated into single cells using Accutase (Innovative Cell Technologies, Inc., Cat# AT104-500), according to the manufacturer’s instructions. An appropriate number of cells were seeded in mTeSR Plus medium supplemented with 10 µM Y-27632 (ROCK inhibitor; MilliporeSigma, Cat# 5092280001) and cultured for 24 hours. The following day, the medium was replaced to remove the ROCK inhibitor, and transfection was performed. Silencer Select siRNAs targeting human *CFDP1* or *SRCAP* (Thermo Fisher Scientific, Cat# 4392420) were used for gene knockdown. A non-targeting negative control siRNA (Scramble siRNA; Thermo Fisher Scientific, Cat# 4390846), with no significant sequence similarity to any known human gene, was used to assess knockdown specificity and serve as a baseline for functional analysis. Transfections were performed using Lipofectamine RNAiMAX Reagent (Invitrogen, Cat# 13778075) at a final siRNA concentration of 100 nM, following the manufacturer’s protocol. Cells were incubated with siRNA for 48 hours prior to downstream analyses.

### qRT-PCR analysis

Total RNA was extracted using TRIzol Reagent (Invitrogen, Cat# 15596026) according to the manufacturer’s instructions. cDNA was synthesized using the SuperScript™ IV First-Strand Synthesis System (Invitrogen, Cat# 18091200). Quantitative real-time PCR (qRT-PCR) was performed using KAPA SYBR® FAST qPCR Master Mix (Sigma, Cat# KK4601) on a LightCycler 480 Real-Time PCR System (Roche). The qRT-PCR primer sequences used are listed in **Supplementary Table 2**.

### Chromatin extraction and western blot

Chromatin extraction was performed as described in our previous publication (*46*). Briefly, 10⁶–10⁷ cells were washed with ice-cold PBS supplemented with 5 mM sodium and lysed in a buffer containing 0.25 M sucrose, 3 mM CaCl₂, 1 mM Tris (pH 8.0), and 0.5% NP-40. Nuclei were pelleted by centrifugation at 3,900 rpm for 5 minutes at 4 °C and washed with either low- or high-salt wash buffer, and centrifuged again under the same conditions. The nuclear pellet was resuspended in extraction buffer (0.5 M HCl, 10% glycerol, 0.1 M 2-mercaptoethylamine HCl), incubated on ice for 30 minutes, and centrifuged at 13,000 rpm for 5 minutes at 4 °C. The supernatant containing histones was precipitated with acetone overnight at −20 °C and centrifuged for collection. Histone samples were analyzed by western blot using rabbit anti-H2A.Z (Abcam, ab4174) and rabbit anti-H3 (Proteintech, 17168-1-AP) antibodies.

### CUT&RUN

#### Library Preparation

CUT&RUN was performed using the CUTANA® ChIC/CUT&RUN Kit (EpiCypher, Cat#14–1048) according to the manufacturer’s instructions. Briefly, hiPSCs were harvested using Versene (Thermo Scientific, Cat# 15040-066), and cell viability was assessed using Trypan Blue staining with the Countess 3 Automated Cell Counter (Invitrogen), ensuring that only samples with >95% viable cells were processed. A total of 0.5 × 10⁶ cells were used per sample. On Day 1, Concanavalin A magnetic beads were activated with Bead Activation Buffer and incubated with cells to facilitate binding. After adding the K-MetStat Panel, cells were then incubated overnight at 4 °C with gentle rocking in Antibody Buffer containing primary antibody of H3K4me3 (EpiCypher, Cat#13-0060), H3K27me3 (EpiCypher, Cat#13-0055), H2A.Z (Abcam, Cat#ab4174) or IgG control (EpiCypher, Cat#13-0042). On Day 2, bead-bound cells were washed twice with cold Cell Permeabilization Buffer, followed by incubation with pAG-MNase at room temperature for 15 minutes. After two additional washes with cold Cell Permeabilization Buffer, calcium was added to activate MNase, and targeted chromatin digestion was carried out at 4 °C for 2 hours. Digestion was stopped by adding Stop Buffer supplemented with E. coli spike-in DNA, and chromatin fragments were released by incubating samples at 37 °C for 10 minutes. DNA-containing supernatants were collected following magnetic separation. DNA was purified using SPRIselect beads, washed twice with 85% ethanol, and eluted in 0.1X TE Buffer. Eluted DNA was analyzed using the FEMTO Pulse system (Advanced Analytical) to assess fragment size distribution prior to library preparation. Two independent biological replicates were performed with minor protocol adjustments to exclude phenotypes resulting from technical variability. In the first round of experiments, 5× MNase and 2 μL of the K-MetStat Panel were used, with results presented in **Supplementary Fig. 4**. In the second round, 3× MNase and 3 μL of the K-MetStat Panel were applied, and the results are shown in **Fig. 4**.

#### CUT&RUN analysis

CUT&RUN libraries were prepared from CUT&RUN DNA using the NEBNext® Ultra™ II DNA Library Prep Kit for Illumina (New England Biolabs, Cat#E7645L), and sequenced using paired-end 50 bp reads on the Singular Genomics G4 sequencer (Singular Genomics). Fastq files were processed using the nf-core/cutandrun pipeline v3.1 using nextflow v. 23.04.1 in a singularity environment (*47*). Briefly, fastq files stemming from each sequencing library were merged and reads were trimmed using TrimGalore and aligned with bowtie2 (minimum qscore of 20 and bin size of 50bps were used for normalization) against hg38. In parallel, reads stemming from the SNAP-CUTANA chromatin-mark-specific spiked-in histones were quantified by counting perfectly-matching reads. Duplicate genomic reads were marked using Picard [https://broadinstitute.github.io/picard/] and bedGraph files were generated using bedtools (*48*), using either the E. coli spike-in data or the chromatin-specific SNAP-Cutana spike-ins. Most downstream analyses were done based on SNAP-cutana normalized bedGraph files. Spike-in normalized files were used to generate bigWig and peak calls using SEACR (*49*) in stringent mode, without normalization (as the files had been previously normalized using the SNAP-Cutana spike-ins). More specifically, each histone or H2AZ mark file was run using both control or treatment IgG controls and the resulting peak files were merged using bedtools merge (flags -c 4,5,6 -o collapse). Similarly, condition-specific consensus peaks were generated from concatenating and merging peak files from each replicate within each condition. Sample-specific coverages were calculated for each control consensus peak using bedtools multicov. The resulting count matrices were filtered for peaks with at least 50 reads across the six samples and imported in the R statistical environment (v 4.2.0) and processed using DESeq2 (*50*).After building a standard DESeq2 quantification object, size factors were substituted with spike-in derived size factors obtained from SNAP-CUTANA counts for the whole spike-in panel in each sample, followed by a re-estimation of dispersion and recomputation of the negative binomial Wald test. Finally, differential binding was evaluated using a normal model comparing siCFDP1 and siSCR conditions. Peaks displaying a false-discovery rate <5% were retrieved and subjected to GREAT analysis (v. 4.0.4) (*51, 52*), using the following association rule: basal+extension: 5000 bp upstream, 1000 bp downstream, 1000000 bp max extension, curated regulatory domains included.

## ACKNOWLEDGEMENTS

We thank members of the Peterson and Boyer Labs for their constructive discourse. We thank Jacques Cote for providing cDNAs of several SRCAP subunits and Shintaro Iwashita for providing CFDP1 cDNA in the initial stage of this work.

## FUNDING

This work was supported by grants from the NIH to C.L.P. (R35GM122519) and L.A.B. (R01HL140471), and the American Heart Association and the NIH to S.W. (16SDG31400009, R03HD095088, R01GM134130),

## CONFLICT OF INTEREST

The authors declare no competing interests.

## AUTHOR CONTRIBUTION

N.M. and S.W. performed protein purifications and biochemical analyses.

D.Y. performed chromatin extraction and CUT&RUN of H2A.Z in human iPSCs.

V.L.B. and S.S.L. conducted the bioinformatic analysis.

C.L.P., L.A.B., and S.W. were responsible for project overview and data interpretations.

All authors assisted in manuscript preparation.

## DATA AVAILABILITY

The data are available from the corresponding author upon reasonable request.

## Notes

### Competing Interest Statement

The authors have declared no competing interest.

