## Supplemental Tables and Figures for "CFDP1 is required for histone variant H2A.Z deposition by the human SRCAP chromatin remodeling complex"

### Supplementary table 1

**Supplementary Table 1. SRCAP-C subunit stoichiometries determined by mass spectrometry.** Data are presented as means and standard deviations from six technical replicates and two independent preparations.

| | WT (High-salt) | WT (Low-salt) |
|---|---|---|
| SRCAP | 1.000 | 1.000 |
| RUVBL1 | 3.680 ± 0.392 | 2.813 ± 0.915 |
| RUVBL2 | 2.455 ± 0.188 | 2.029 ± 0.439 |
| DMAP1 | 0.801 ± 0.241 | 0.723 ± 0.069 |
| Actin | 1.277 ± 0.269 | 0.838 ± 0.154 |
| BAF53a | 0.408 ± 0.090 | 0.423 ± 0.108 |
| GAS41 | 0.350 ± 0.051 | 0.600 ± 0.242 |
| YL1 | 0.963 ± 0.194 | 0.692 ± 0.209 |
| ARP 6 | 0.304 ± 0.073 | 0.342 ± 0.036 |
| ZNHIT1 | 0.481 ± 0.117 | 0.410 ± 0.120 |
| CFDP1 | 0.001 ± 0.001 | 0.059 ± 0.011 |
| BRD8 | 0.001 ± 0.001 | 0.004 ± 0.002 |

### Supplementary table 2

**Supplemental Table 2. List of qPCR primer sequences used in this study**

| | |
|---|---|
| *hCFDP1-F* | *TGCTGGTGAAGAAGTAAGGGTA* |
| *hCFDP1-R* | *TTTCCCCAAAAGGCTGCTCA* |
| *hSRCAP-F* | *GGCTGGTTACCATGTATGAGAAG* |
| *hSRCAP-R* | *CATCACGCTGGTGGGAACAA* |
| *hOct4-F* | *CTTGAATCCCGAATGGAAAGGG* |
| *hOct4-R* | *CCTTCCCAAATAGAACCCCCA* |
| *hSox2-F* | *GCCGAGTGGAAACTTTTGTCG* |
| *hSox2-R* | *GGCAGCGTGTACTTATCCTTCT* |
| *hNanog-F* | *AAGGTCCCGGTCAAGAAACAG* |
| *hNanog-R* | *CTTCTGCGTCACACCATTGC* |
| *hH2A.Z-F* | *GGACGACCAGTCATGGACG* |
| *hH2A.Z-R* | *TGACGAGGGGTAATACGCTTT* |
| *hGata1-F* | *TTGTCAGTAAACGGGCAGGTA* |
| *hGata1-R* | *CTTGCGGTTTCGAGTCTGAAT* |
| *h Brachyury -F* | *CTGGGTACTCCCAATGGGG* |
| *hBrachyury -R* | *GGTTGGAGAATTGTTCCGATGA* |
| *hBmp2-F* | *ACCCGCTGTCTTCTAGCGT* |
| *hBmp2-R* | *TTTCAGGCCGAACATGCTGAG* |
| *hBmp4-F* | *AAAGTCGCCGAGATTCAGGG* |
| *hBmp4-R* | *GACGGCACTCTTGCTAGGC* |
| *hGata3-F* | *GCCCCTCATTAAGCCCAAG* |
| *hGata3-R* | *TTGTGGTGGTCTGACAGTTCG* |
| *hHoxa1-F* | *TCCTGGAATACCCCATACTTAGC* |
| *hHoxa1-R* | *GCACGACTGGAAAGTTGTAATCC* |
| *hHoxa5-F* | *TACGGCTACAATGGCATGGAT* |
| *hHoxa5-R* | *CCGCTGGAGTTGCTTAGGG* |
| *hHoxa6-F* | *TCCCGGACAAGACGTACAC* |
| *hHoxa6-R* | *CGCCACTGAGGTCCTTATCA* |
| *hHoxb2-F* | *CGCCAGGATTCACCTTTCCTT* |
| *hHoxb2-R* | *CCCTGTAGGCTAGGGGAGAG* |
| *hHoxb9-F* | *CCGTCTACCACCCTTACATCC* |
| *hHoxb9-R* | *CGTAGCCGGGTCTTTGATTAG* |
| *hHoxb13-F* | *CCAGTTACCTGGACGTGTCTG* |
| *hHoxb13-R* | *GGACCTGGTGGGTTCTGTTC* |

### Supplementary Figure 1

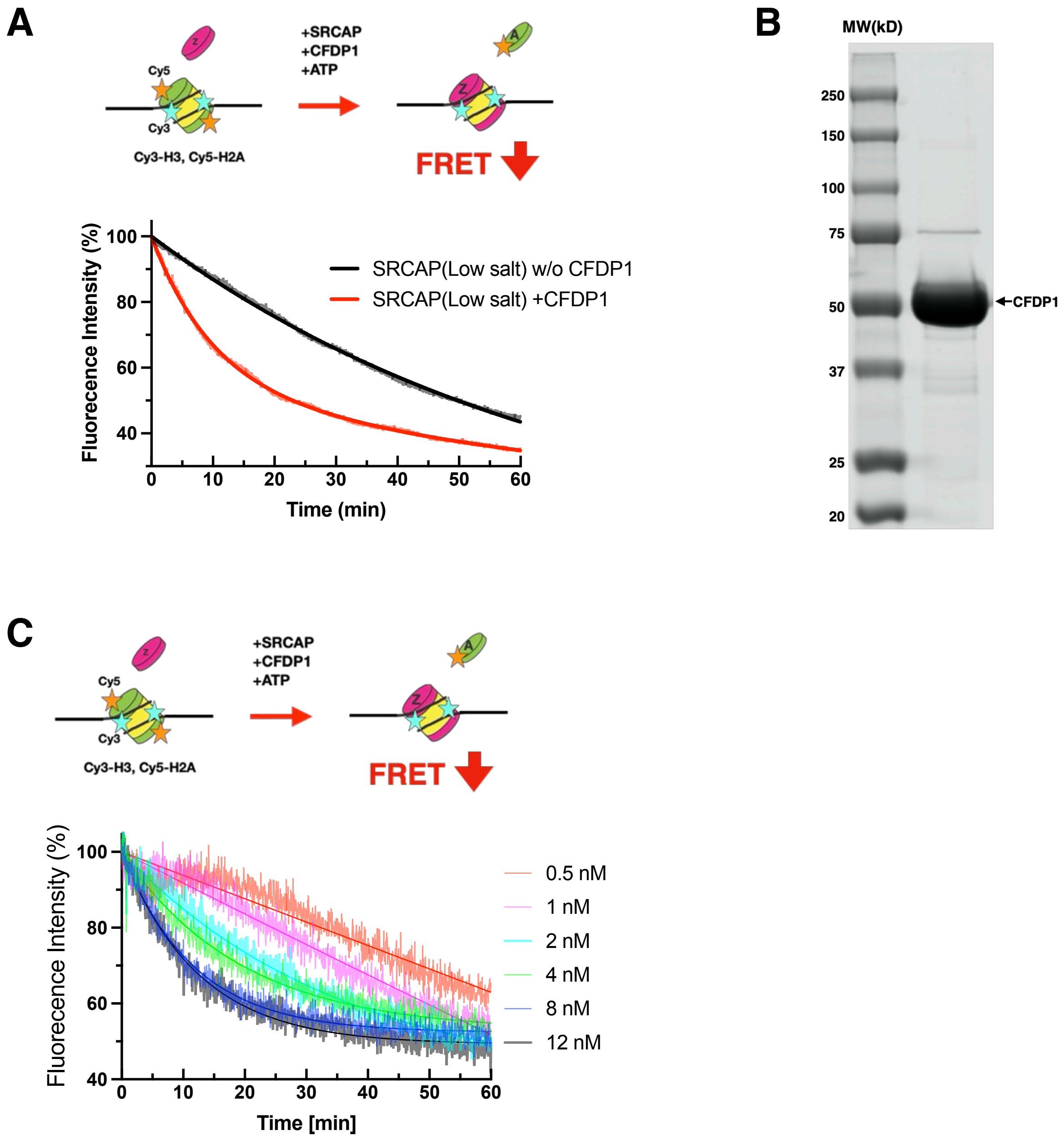

**Supplementary Figure 1.**

(A) Dimer exchange activity of SRCAP-C purified under the low-salt condition. Scheme of FRET-based dimer eviction assay (Top panel). Representative changes in FRET fluorescence intensity over time using a PC1 fluorescent spectrometer (70 μL reaction volume). The assay was performed as described in Fig. 1C.

(B) Coomassie-staining SDS-PAGE of the purified FLAG-tagged CFDP1.

(C) SRCAP-concentration dependence of dimer exchange activity. Scheme of FRET-based dimer eviction assay (Top panel). Representative changes in FRET fluorescence intensity over time using a Tecan plate reader (70 μL reaction volume) (Bottom panel). The assay was performed as described in Fig. 1C.

### Supplementary Figure 2

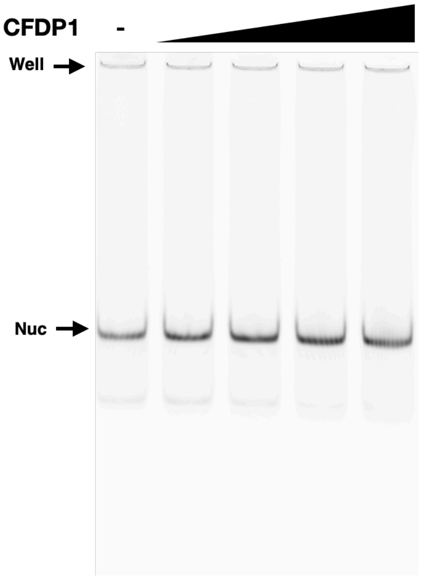

**Supplementary Figure 2.**

Representative native gel of EMSA for the nucleosome binding of CFDP1. 7 nM mononucleosomes were incubated with CFDP1 (0.5, 1, 2, and 4 μM) and subjected to 4% Native-PAGE.

### Supplementary Figure 3

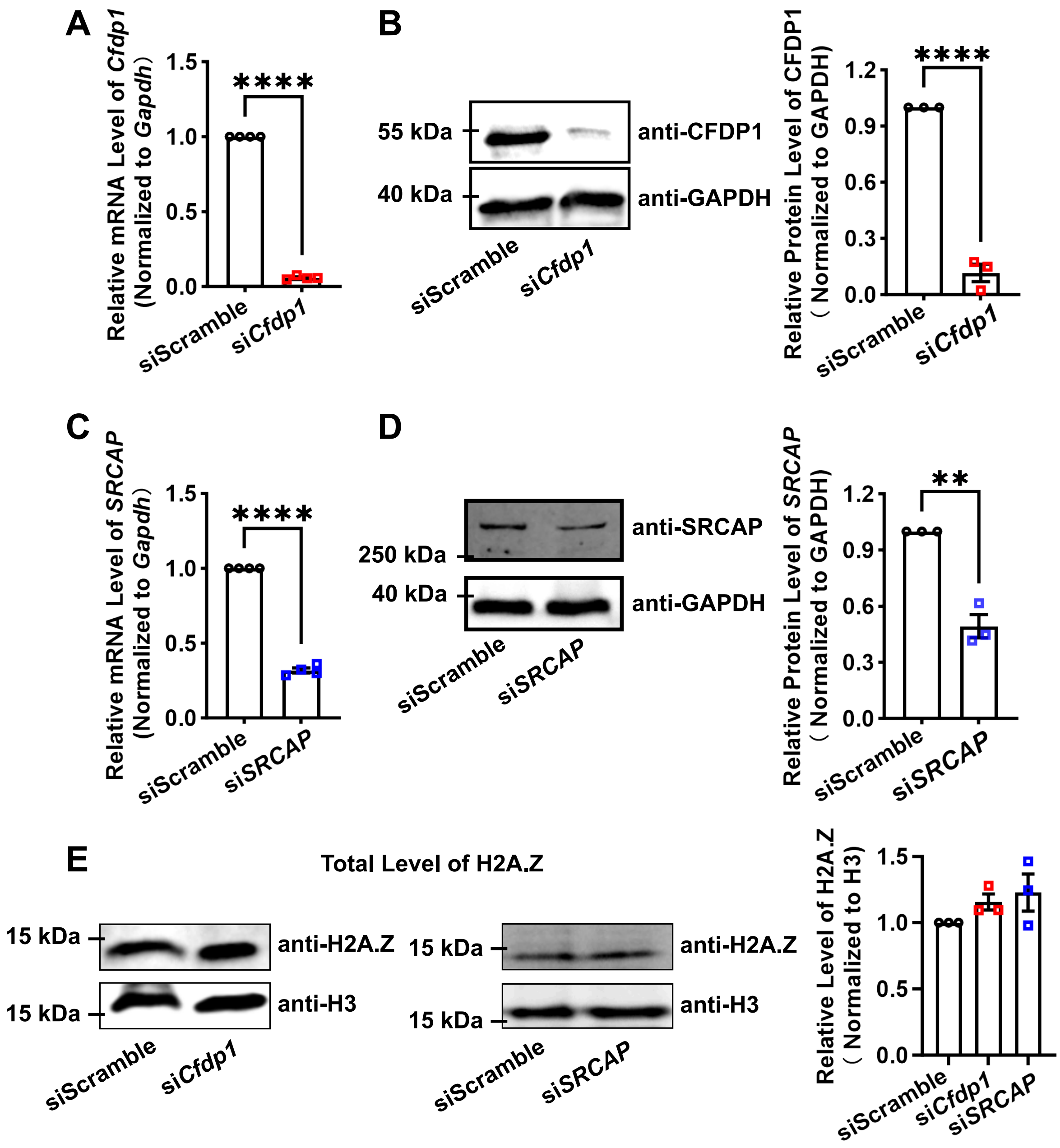

**Supplementary Figure 3**.

Depletion of *CFDP1* or *SRCAP* has minimal effect on total H2A.Z levels in hiPSCs.

(A–D) Validation of siRNA-mediated knockdown efficiency for *CFDP1* (A–B) and *SRCAP* (C–D) at both mRNA and protein levels by qRT-PCR (A, C) and Western blot (B, D), respectively. *GAPDH* was used as a normalization control. Data are presented relative to expression levels in siScramble cells. $N \geq 3$; error bars represent 2 standard deviations from the mean. $^{**}P < 0.01$, $^{****}P < 0.0001$ versus siScramble. (E) Examination of total H2A.Z levels revealed no significant difference between si*CFDP1* and siScramble cells. Histone H3 was used as a loading control. $N=3$; error bars represent 2 standard deviations from the mean.

### Supplementary Figure 4

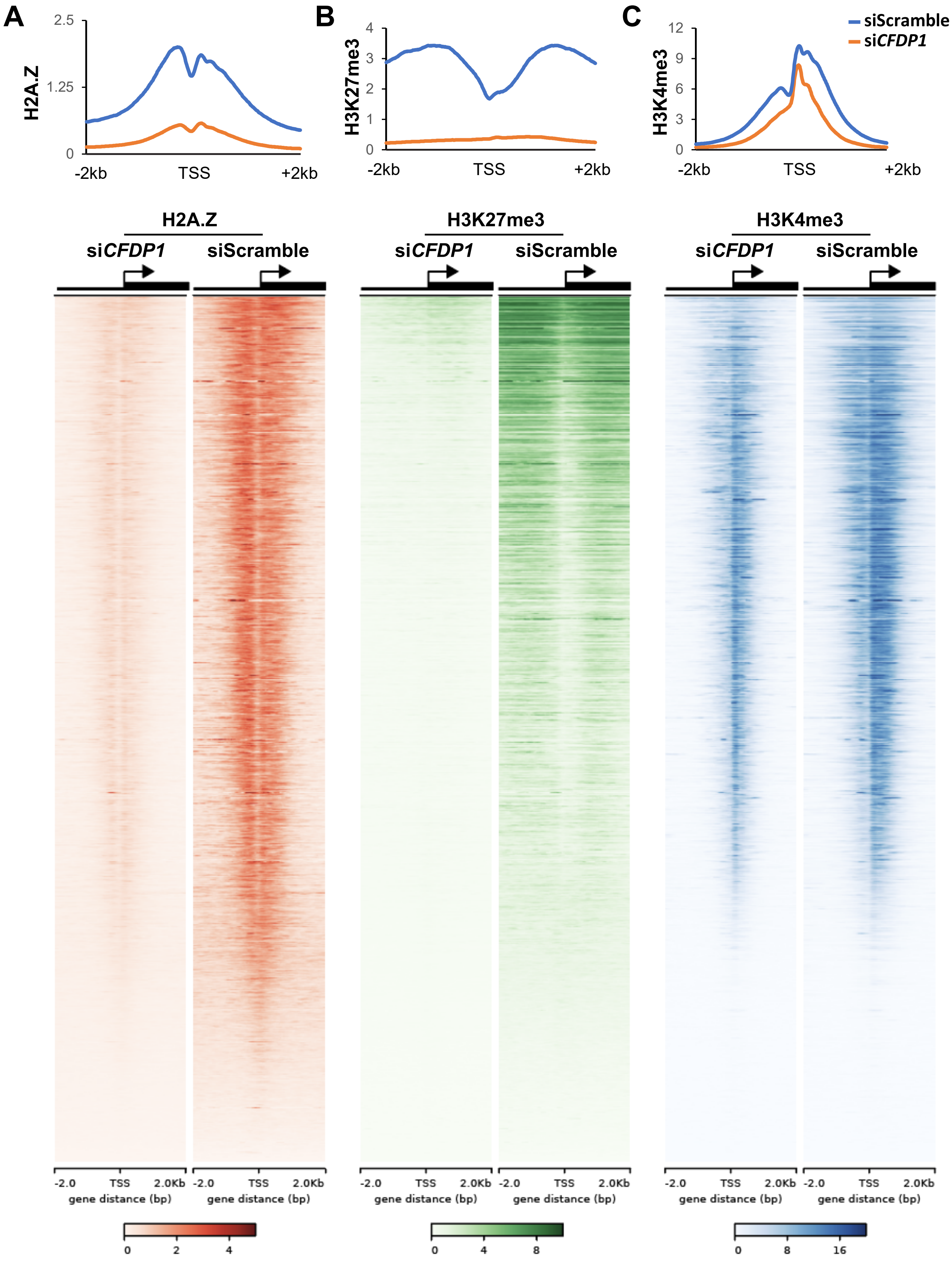

**Supplement Figure 4.**

Loss of *CFDP1* reduces H2A.Z, H3K27me3, and H3K4me3 enrichment in hiPSCs (independent biological replicates).

(A–C) Average enrichment profiles and density maps of H2A.Z (A), H3K27me3 (B), and H3K4me3 (C) within ±2 kb of transcription start sites (TSSs), showing a global decrease in *CFDP1*-deficient cells compared to siScramble controls. In density maps, genes are ordered by H2A.Z signal, with enrichment shown across a ±2 kb window around the TSS.
